## Supplemental Information for "Integration of mouse ovary morphogenesis with developmental dynamics of the oviduct, ovarian ligaments, and rete ovarii"

#### Supplemental Figures

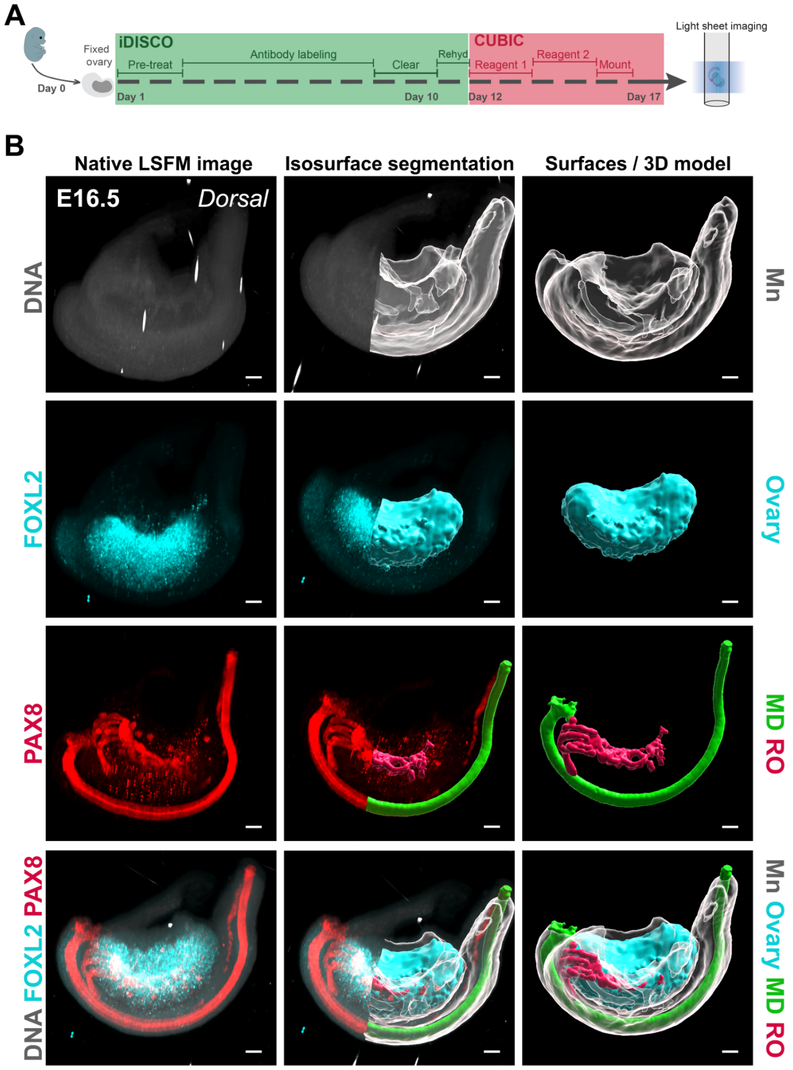

**Figure S1. Approach to generate 3D models of the developing ovary.** (**A**) Schematic depicting the basic pipeline for producing 3D images of the fetal mouse ovary using iDISCO+CUBIC tissue clearing and lightsheet microscopy. (**B**) Approach for generating 3D models in Imaris software using isosurface segmentation of lightsheet images for each of the main markers used in this study The left column shows the 3D *maximum intensity projection* view of the raw lightsheet Z-stack of an ovary/mesonephros complex E16.5 immunostained for FOXL2 (*cyan*) and PAX8 (*red*), and counterstained with Hoechst nuclear dye (*grayscale*); the middle column shows the process of isosurface segmentation in Imaris and the right column shows the final 3D surfaces used in the figures throughout the manuscript. MD, Mullerian duct; RO, rete ovarii; Mn, mesonephros. Scale bars, 100µm.

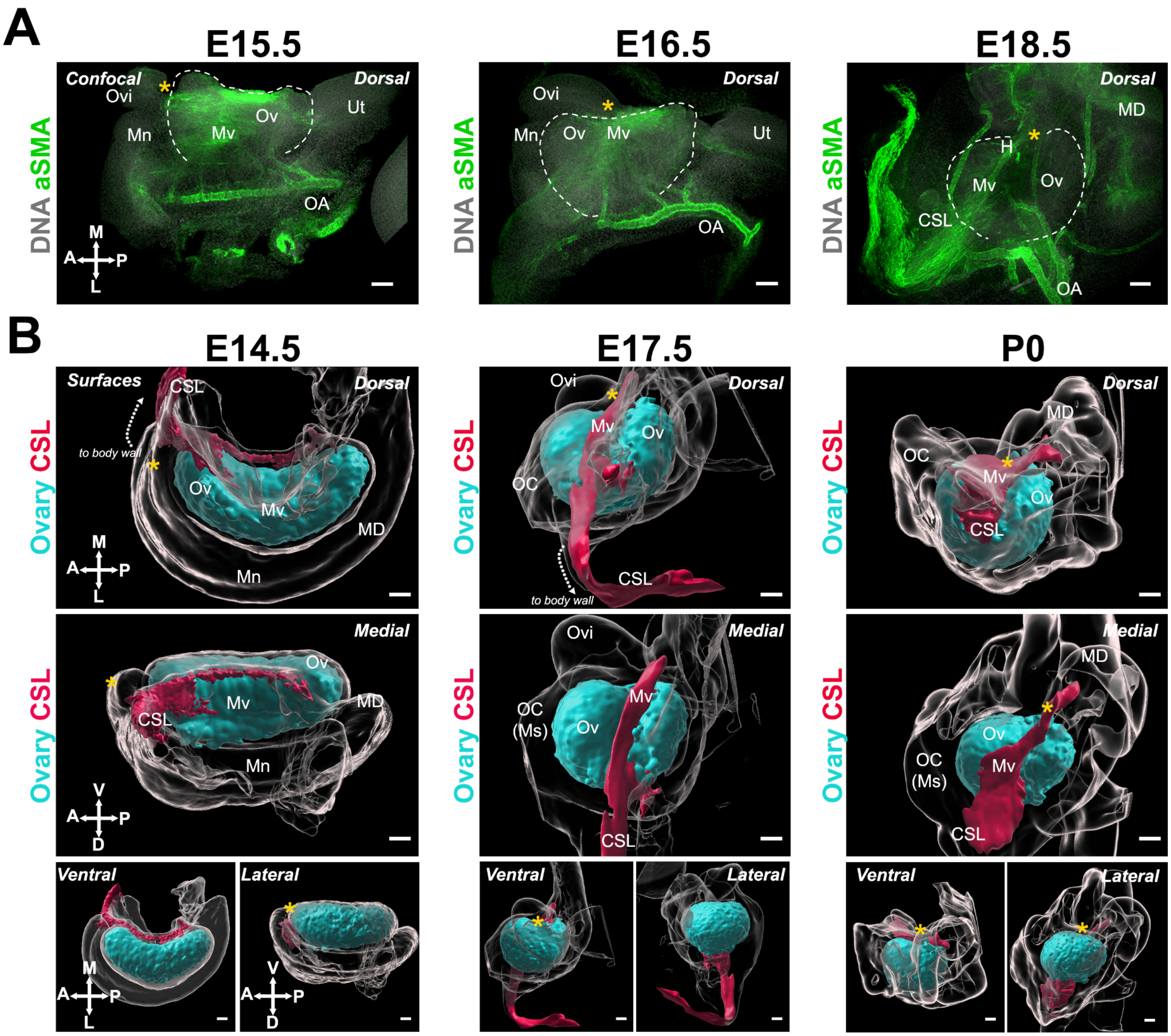

**Figure S2. The mesovarium and cranial suspensory ligament tether the ovary to the rest of the urogenital complex.** (**A**) Confocal images of whole ovary/mesonephros complexes at E15.5, E16.5 and E18.5 immunostained for aSMA (*green*), and counterstained with Hoechst nuclear dye (*grayscale*). Samples in the top two rows were imaged from the dorsal side and samples in the bottom row from the medial side. (**B**) 3D models generated by isosurface segmentation of lightsheet images taken of whole ovaries at E14.5, E17.5 and P0 immunostained for FOXL2 (*cyan*) and TNC (E14.5, *red*) or labelled with AF647-Hydrazine to reveal Elastin expression (E17.5, P0**,** *red*). All samples were counterstained with Hoechst nuclear dye (*grayscale*). Top panels represent the dorsal view, middle panels represent the medial view, and small bottom panels illustrate ventral and lateral views of the same ovary. White dashed arrows show the direction to the body wall. Yellow asterisks indicate the location of the infundibulum of the presumptive oviduct for reference. Compasses on the bottom left of each panel indicate the orientation of the ovary for the entire row: V, ventral; L, lateral; D, dorsal; M, medial. CSL, cranial suspensory ligament; H, hilum; MD, Müllerian duct; Mm, mesometrium; Mn, mesonephros; Ms, mesosalpinx; Mv, mesovarium; OA, ovarian artery; OC, ovarian capsule; Ov, ovary; Ovi, oviduct; Ut, uterus. Scale bars, 100μm.

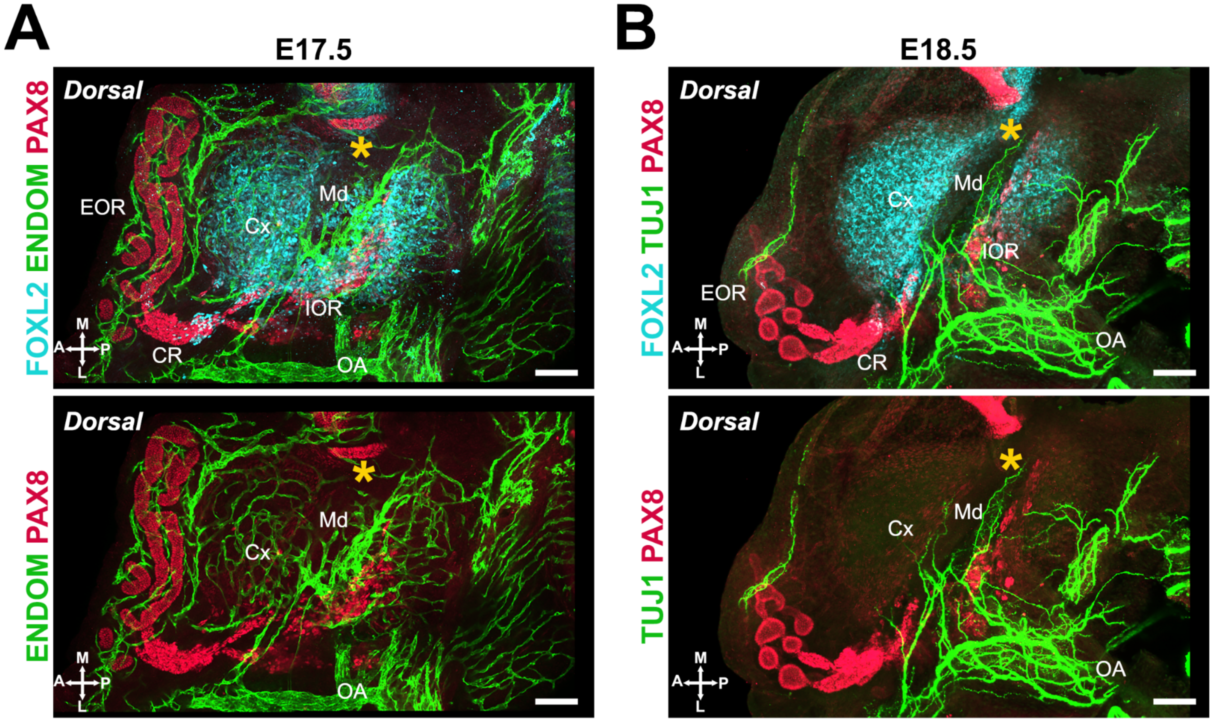

**Figure S3. The IOR and CR coincide with the point of entry of vasculature and innervation into the ovary.** (**A**) Maximum intensity projections from confocal Z-stacks of the dorsal side of the whole ovary/mesonephros complex at E17.5 immunostained for ENDOMUCIN (*green*) and PAX8 (*red*). (**B**) Maximum intensity projections from confocal Z-stacks of a whole ovary/mesonephros complex at E18.5 immunostained for TUJ1 (*green*) and PAX8 (*red*), imaged from the dorsal side. Yellow asterisks indicate the location of the infundibulum of the presumptive oviduct for reference. CR, connecting rete; Cx, cortex; EOR, extraovarian rete; IOR, intraovarian rete; Md, medulla; OA, ovarian artery. Compasses on the bottom left of each panel indicate the orientation of the ovary: V, ventral; L, lateral; D, dorsal; M, medial. Scale bars, 100μm.

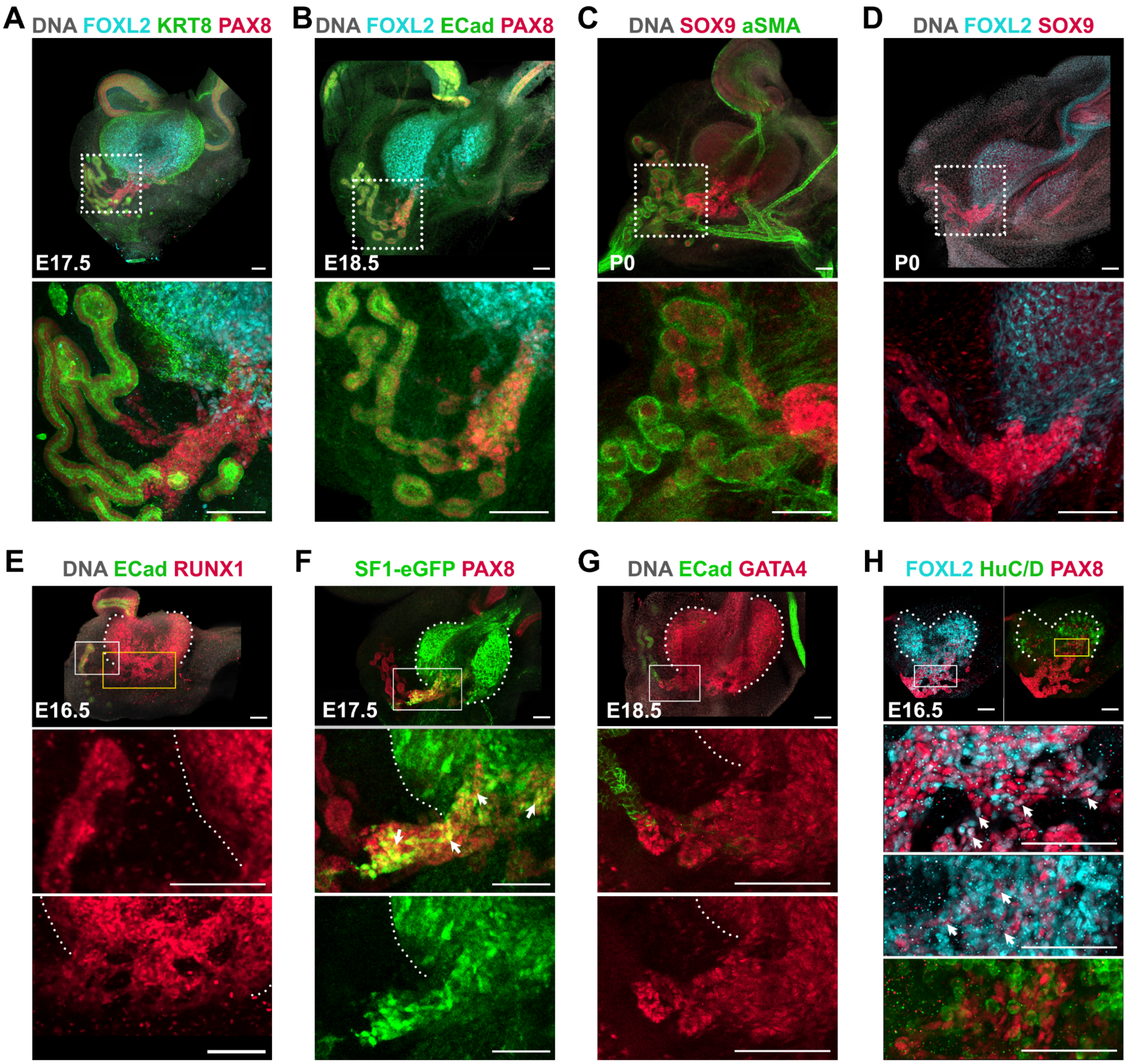

**Figure S4. Expression of known epithelial and gonadal markers in the different regions of the rete ovarii.** (**A-D**) Maximum intensity projections from confocal Z-stacks of whole ovary/mesonephros complexes at E17.5, E18.5 or P0, imaged from the dorsal side. Images in the bottom row are close-ups of the areas outlined in the top row. (**A**) immunostaining for FOXL2 (*cyan*), PAX8 (*red*) and KERATIN-8 (KRT8, *green*). (**B**) immunostaining for FOXL2 (*cyan*), PAX8 (*red*) and E-CADHERIN (ECad, *green*). (**C**) immunostaining for SOX9 (*cyan*) and aSMA (*red*). (**D**) immunostaining for FOXL2 (*cyan*) and SOX9 (*red*). (**E-H**) Maximum intensity projections from confocal Z-stacks of whole ovary/mesonephros complexes from wild-type (**E, G, H**) or Sf1-eGFP (**F**) embryos at E16.5 (**E, H**), E17.5 (**F**), or E18.5 (**G**), imaged from the dorsal side. (**E**) immunostaining for RUNX1 (*red*) and E-CADHERIN (ECad, *green*). (**F**) immunostaining E17.5 Sf1-eGFP for GFP (*green*) and PAX8 (*red*). (**G**) immunostaining for ECad (*cyan*) and GATA4 (*red*). (**H**) immunostaining for FOXL2 (*cyan*), PAX8 (*red*) and HuC/D (*green*). White arrows point to PAX8+ cells that are also labeled with gonadal markers. Images in the bottom rows are close-ups of the areas outlined in the top row. Yellow rectangles in **E** and **H** outline the regions blown up in the lowest row. Scale bars, 100µm.

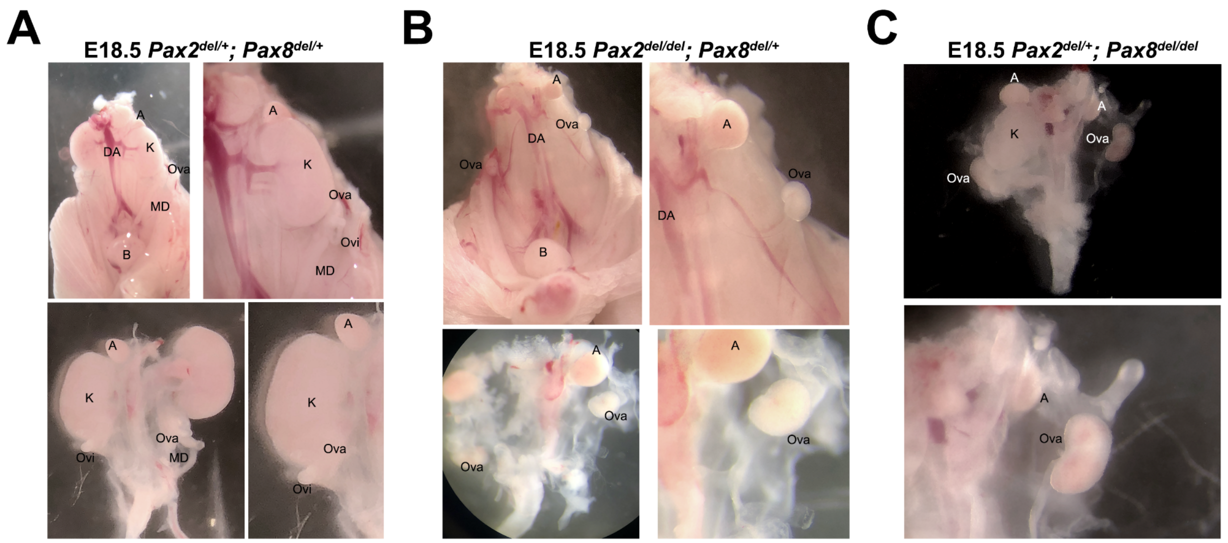

**Figure S5. Gross morphology of *Pax2*$^{del}$; *Pax8*$^{del}$ fetuses.** (**A-B**) Macrophotography of the urogenital complex *in situ* (*top row*) or isolated (*bottom row*) in *Pax2*$^{del/+}$*; Pax8*$^{del/+}$ (**A**) or *Pax2*$^{del/del}$*; Pax8*$^{del/+}$ (**B**) fetuses at E18.5. (**C**) Macrophotography of the isolated urogenital complex from a *Pax2*$^{del/+}$*; Pax8*$^{del/del}$ fetus at E18.5. Note the absence of kidneys (K) in **B** and **C**, and the resulting shorter distance between ovary (ova) and adrenal (A) compared to **A**. A, adrenal; B, bladder; DA, dorsal aorta; K, kidney; MD, Müllerian duct; Ova, ovary; Ovi, oviduct.

### Supplemental Tables

**Supplemental Table 1.** Ovarian and Müllerian duct phenotypes in *Pax2* and *Pax8* allelic series

| | | ***P2+/+ P8+/+*** | ***P2d/+ P8+/+*** | ***P2d/d P8+/+*** | ***P2+/+ P8d/+*** | ***P2+/+ P8d/d*** | ***P2d/+ P8d/+*** | ***P2d/d P8d/+*** | ***P2d/+ P8d/d*** |
|---|---|---|---|---|---|---|---|---|---|
| **Müllerian duct** | **Oviduct** | Intact | Perturbed | Perturbed | Intact | Perturbed | Perturbed | Absent | Absent |
| | **Infundibulum** | Intact | Intact | Intact | Intact | Intact | Intact | Intact | Intact |
| **Rete ovarii** | **IOR** | Intact | Intact | Intact | Absent | Absent | Perturbed | Absent | Absent |
| | **CR** | Intact | Intact | Intact | Perturbed | Absent | Perturbed | Perturbed | Absent |
| | **EOR** | Intact | Intact | Absent | Intact | Absent | Absent | Absent | Absent |
| **Ovary morphogenesis** | **Folding** | Intact | Intact | Intact | Intact | Intact | Intact | Intact | Intact |
| | **Encapsulation** | Intact | Intact | Intact | Intact | Absent | Absent | Absent | Absent |

**Legend**

| Intact | Perturbed | Absent |
|---|---|---|

**Supplemental Table 2.** PCR Primers used in this study

| Allele | Forward Primer | Reverse Primer |
|---|---|---|
| *Pax2 wild-type* | AAAGTGAGGGAAGCGTAGAGAAG | ACCATAGACATTAGAGGTGCAGA |
| *Pax2del* | TGACTTTTGCAGTCCAGAGTCTCCC | ACCATAGACATTAGAGGTGCAGA |
| *Pax8 wild-type* | GAAAGTTCGAGGGAAGGGAGATC | CAGTTCTTTCAGTGGTCCCTCC |
| *Pax8del* | GAAAGTTCGAGGGAAGGGAGATC | GACCAAGTAGTCAATGGGGGCC |
| *Sf1:eGfp* | CACCATCTTCTTCAAGGACGAC | GTCACGAACTCCAGCAGGACC |
| *Sry* | GTGTCTCAAAGCCTGCTCTTC | CATGTACTGCTAGCAGCTATC |
| *Myogenin (internal control)* | TTACGTCCATCGTGGACAGCAT | TGGGCTGGGTGTTAGTCTTAT |

**Supplemental Table 3.** Primary antibodies used in this study

| Primary Antibody | Host Species | Dilution | Source | Product # |
|---|---|---|---|---|
| AMH/MIS | Goat | 1:500 | Santa Cruz Biotechnology | sc-6886 (discontinued) |
| aSMA (Cy3 conjugated) | Mouse | 1:1000 | Sigma | C6198 |
| aSMA (FITC conjugated) | Mouse | 1:500 | Sigma | F3777 |
| E-Cadherin | Rat | 1:500 | Zymed (Thermo Fischer) | 13-1900 |
| ENDOMUCIN | Rat | 1:250 | Santa Cruz Biotechnology | sc-65495 |
| FOXL2 | Goat | 1:250 | Novus Biologicals | NB100-1277 |
| GATA4 | Goat | 1:250 | Santa Cruz Biotechnology | sc-1237 (discontinued) |
| GFP | Chicken | 1:1000 | Abcam | ab13970 |
| HuC/D | Human | 1:10000 | Gift from V. Lennon (Mayo Clinic) | N/A |
| KRT8 | Rat | 1:250 | DSHB | TROMA-I |
| PAX8 | Rabbit | 1:500 | Proteintech | 10336-1-AP |
| RUNX1 | Rabbit | 1:500 | Abcam | ab92336 |
| SOX9 | Rabbit | 1:1000 | Millipore | AB5535 |
| TNC | Rabbit | 1:250 | Gift from H. Erickson (Duke University) | N/A |
| TUJ1 | Rabbit | 1:1000 | Abcam | ab18207 |

**Supplemental Table 4.** Secondary antibodies used in this study

| Secondary Antibody | Dilution | Source | Product # |
|---|---|---|---|
| AF647 Donkey anti-**Rabbit** | 1:500 | Jackson ImmunoResearch | 711-605-152 |
| Cy3 Donkey anti-**Goat** | 1:500 | Jackson ImmunoResearch | 705-165-147 |
| Cy3 Donkey anti-**Chicken** | 1:500 | Jackson ImmunoResearch | 703-165-155 |
| AF488 Donkey anti-**Chicken** | 1:500 | Jackson ImmunoResearch | 703-545-155 |
| AF488 Donkey anti-**Human** | 1:500 | Jackson ImmunoResearch | 709-545-149 |
| AF488 Donkey anti-**Rat** | 1:500 | Life Technologies | A-21208 |
| Cy3 Donkey anti-**Rat** | 1:500 | Jackson ImmunoResearch | 712-165-150 |
| CF647-hydrazide probe to label Elastin | 1:500 | Millipore Sigma | SCJ4600046 |

#### Supplemental movie legends

**Movie S1. 3D model of an XX embryo at E14.5.** Imaris isosurface segmentation tool was used to generate 3D models of the whole embryo volume (*grayscale*, based on background staining), the kidneys (*green*, based on PAX8 immunolabeling), the reproductive tract epithelia (*red*, based on PAX8 labelling), and the ovary (*cyan*, based on FOXL2 immunolabeling). The video shows rotation and close-ups of the sample to view different aspects of the complexes in their native context.

**Movie S2. Imaris workflow.** The video shows the 3D rendering of native lightsheet images of an E14.5 ovary mesonephros complex labelled with FOXL2 to visualize the gonad (*cyan*), PAX8 to visualize the reproductive ducts (*red*), and Hoechst nuclear dye to visualize the whole complex (*grayscale*). The next frames show the transition from native data to 3D models using isosurface segmentation in Imaris software and rotation of the sample to capture different 3D views of the ovary and mesonephros in their native conformation.
